# Diffusion assessment through image processing: beyond the point-source paradigm

**DOI:** 10.1101/2020.05.27.118588

**Authors:** Eugene B. Postnikov, Anton A. Namykin, Oxana V. Semyachkina-Glushkovskaya, Dmitry E. Postnov

## Abstract

The quantification of transport processes of different substances in the brain’s parenchyma is important in the context of understanding brain functioning. Most of the currently used methods for assessment of the effective diffusion coefficient rely on the point-source paradigm.

We propose a method for the quantitative characterization of the diffusion process in the brain’s parenchyma using a set of images recorded in the experiment during the spreading of a fluorescent dye. Our method exploits the frame-wise comparison of experimental data with a set of images that would be observed for an ideal diffusion process within the same topology. We obtain this reference set of images using blurring the image with an appropriate kernel function, and the degree of such blurring correlates with the spreading process of a dye. We demonstrate the applicability of the proposed method using (i) the simulated surrogate data, (ii) the set of experimentally recorded fluorescent images of the isolated event of blood-brain barrier (BBB) opening, and (iii) the images of massive multi-source spreading of fluorescent dye.

## Introduction

The problem of understanding and quantitative assessment of transport processes in the brain’s parenchyma was termed as “the final frontier of neuroscience” [1]. It is important in the context of different tasks related to the understanding of brain functioning [2, 3]. The recent example is the active discussion on mechanisms of brain clearance from metabolic products, e.g. amyloid-beta associated with the risk of Alzheimer’s disease [4]. In this context, a so-called glymphatic mechanism has been proposed [5, 6], according to which a directed flow of fluid through the parenchyma makes an important contribution to this process but this hypothesis is disputed [7–9], and the diffusion is considered as the major process that provides both the delivery of useful substances and the removal of waste products.

It should be noted, however, that the spreading of substances in the brain’s parenchyma is a more complex process than the classical diffusion in continuous homogeneous media which is relevant to a variety of problems of physics and physical chemistry.

First of all, instead of a continuous medium, substances spread through the extracellular space (ECS), which forms 10-30% of the total parenchyma’s volume and has a complex shape. As a first approximation, it can be treated as a porous medium, for which the effective diffusion coefficient may be introduced [10, 11]. However, the diffusion through the intercellular space is complicated by additional effects such as a temporary binding of walking molecules to receptors of the cell membranes, interactions with large molecular structures that present in the ECS, or temporary trapping in ECS dead-end segments, see [12] for a comprehensive review. Due to the cumulative effect of such factors, it is almost impossible to predict the effective diffusion coefficient theoretically, therefore, one should rely on experimental methods for its assessment.

Currently, there exist several methods aimed at determining the effective diffusion coefficient, one of the most advanced of which is based on the transfer of microscopic amounts of radioactive molecules into the brain parenchyma by ionophoresis with the subsequent tracking and fitting their further distribution. It should be noted that this method (as well as a number of simpler ones) implies the assumption of the so-called “point-source paradigm” [1, 13, 14], when the marker is inserted by a micropipette exactly in the point, and the marker itself is not very high-molecular. Respectively, this approach works well in vitro with tissue slices and produces a recorded marker distribution close to the Gaussian, which can be compared with the Gaussian distribution recorded in a model uniform medium (the dilute agarose gel). Further, the square root of the ratio of diffusivities provides an estimate for the tortuosity of the ECS in the brain tissue.

Note, there are issues in the application of this and similar methods, especially *in vivo*: Using a syringe needle is traumatic and can create a local tumor that changes the properties of the surrounding extracellular space, the bolus of a substance introduced into the tissue can cause its local non-diffusion transport connected with its slow constrained relaxation, etc. [15].

Also, the point source paradigm is not the best assumption if the substances (usually fluorescent high-molecular compounds, e.g. dextrans) passed through the blood-brain barrier (BBB) as a result of targeted exposure to sound, ultrasound, or laser irradiation [16, 17]. In this case, the elementary sources are the capillary segments, and with respect to the characteristic distance between capillaries (tens of microns), they cannot be considered a point. In addition, many such non-point sources can be distributed over the field of view.

To make quantitative measurements, the measurements of the markers concentration intensity are usually restricted to individual trials and small regions having a simple geometrical shape. The typical choice is a short thin linear region normal to a blood vessel’s boundary that reduces the data processing to one-dimensional problem [18, 19]. This allows one to quantify the influx and spread situation in terms of the diffusive distance, its growth with time [20], and the respective diffusion coefficient. However, these results are weakly applicable to the global characterization of images covering wide regions with the well-developed part of the vascular network having multiple sources of leakage, see the discussion in [21].

At the same time, there are diffusion related approaches in other fields. The determination of intrinsic features of complex patterns (e.g. deformable bodies, graphs, and network structures) can be based on the properties of diffusion on such structures by means of the so-called “diffusion map” [22, 23]. This method exploits the metrical distance between the probability distribution densities taken at the same moment for two distinct points, which can be reached by the diffusion process started from one origin point. This method not only takes into account a variety of available walking paths on a complex structure but also allows a reduced description of the pattern via a small set of leading eigenvalues of the respective diffusion (Fokker-Planck) operators [24]. Moreover, it is proven that the diffusion distance based on the diffusion map eigenfunction/eigenvalue representation does not depend asymptotically on the origin point location for the diffusion process. Although this approach is widespread now in network science, it was not applied yet to the problem of brain fluid spread (rare exceptions are [25, 26]) due to two possible reasons: the original diffusion map method initially adjusted to study of discrete graphs, and imaging data for a continual spread cannot be uniquely characterized by the Euclidean distance between images with a different tool- and condition-dependent intensities of pixels. The latter reason restricts the use of such a method by the segmentation and compressed description of separate images and does not allow a warrant quantitative comparison of real medical imaging data and model simulations.

In our work, we propose a method for the quantitative characterization of the diffusion process in the brain’s parenchyma using a set of images recorded in the experiment and not restricted by highly symmetric (point-wise or linear) regions simultaneously with a set of distributions generated by an ideal diffusion process within the same topology. The essence of the proposed approach is that a 2D image of an ideal diffusion process can be obtained via the Gaussian blurring of the image of a distributed source region, and the degree of blurring reflects the evolution over time. On this basis, for each image recorded in the experiment, the spot luminance formed by the spreading fluorescent marker can be correlated with a certain degree of Gaussian blurring of the source image. The evolution of the blurring degree in time gives information about the rate of spreading from the source, which is the desired effective diffusion coefficient.

The quality of fit by blurred images allows one to estimate quantitatively the degree of applicability of the assumption of the pure diffusion nature of the process by a single numerical parameter. Taken together, these outputs provide information about the features of the spreading process and thus help to recover the underlying physiological events.

In the rest of the paper, we describe the details of the proposed method, demonstrate its work on artificially calculated diffusion processes with different source geometries, and then give the results of data processing of the real experiment on the sound-induced BBB opening.

## Materials and methods

### Correlation-based approach

Suppose we have the sequence of images that represent the process to analyze through the distribution of the brightness of the fluorescent dye. The consideration below is based on the assumption that the spreading process has a diffusive character [9, 19] with the initial condition *u*(*x, y*, 0) = *v*(*x, y*), and *σ*^2^ = 4*Dt*, where *D* is the diffusion coefficient (isotropic) and *t* is the time variable. Note that the accuracy of such an assumption can be checked during the processing algorithm, see below. The diffusion equation reads:

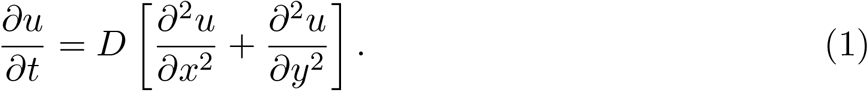

The Cauchy problem solution for Eq. (1), i.e. its Green function, is the Gaussian function. It suggests the most natural kernel for the filtration procedure, which should predict the “true diffusive” blurring of an instant non-point source of the fluorescent marker, which simply plays a role of the initial condition.

In a general case of distributed source *v*(*x, y*), which provide initial influx of the markes into the brain’s space, the blurred picture with the characteristic size (dispersion) *σ* is given by the integral transform

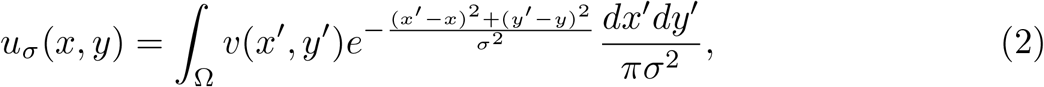

which is written for the whole plane Ω that contains the analyzed image. In the practical situation, when an image is a finite-size recorded frame, Eq. (2) will be replaced by a sum, but we keep its integral form for the convenience of the further mathematical line of reasoning. In addition, one can choose the region of interest in such a way that the analyzed fluorescence region is localized sufficiently far from image boundaries, i.e. one can neglect by boundary effects.

The function *v*(*x, y*) can be determined from the preliminary inspection of the very first of recorded images, where an appropriate mask should be applied to extract 2D indicator function which is equal to 1 in pixels, which outlines the source of the dye.

In order to process each frame in recorded sequence, we propose to compute (2) for a set of ordered dispersion parameters {*σ*_*j*_} with the same *v*(*x, y*). Then, the obtained set if spatial distributions 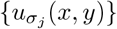 should be compared with the current frame to find *σ*, which provides the best similarity between recorded and blurred images. Note that the discrete pixelized representation of the real processed images makes quite sufficient such discrete quantification to avoid overfitting, which may arise from more sophisticated optimization procedures.

It should be noted that experimental images may valuably differ in their background intensity, range of pixels value magnitude due to the methods of intensity digitalization, etc. For this reason, the Euclidean pairwise distance used in the diffusion map/diffusion distance approach seems to be inappropriate. We replace it by the consideration of the two-dimensional correlation coefficients 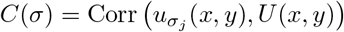 between the experimental images *U* (*x, y*) and the simulated blurred picture since this quantity is invariant with respect to the linear transformation of intensity distributions. In addition, this allows for using an arbitrary magnitude for the initial vascular mask *v*(*x, y*) that simplifies computations.

All practical calculations with image matrices have been done in OCTAVE software, applying the Fast Fourier Transform for computing the convolution integral (2) that follows from the convolution theorem

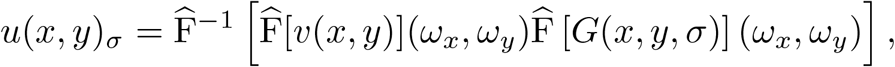

where *G*(*x, y, σ*) denotes the blurring kernel, e.g. the Gaussian kernel in Eq (2), or another one, which can be introduced for more complicated situations, see below.

It should be pointed out that the ideas described and tested above assume an absence of the source acting during any time of the process simulation.

At the same time, the experimentally realistic situation meets a possibility of the continuous leakage of the marker-contained fluid substance from the vessel to the parenchyma. Moreover, this leakage can be non-stationary. In this case, Eq 1 should be supplied with either a time-dependent source term in its right-hand side or a non-zero boundary condition located on the surface of the leakage source. In particular, the simplest case of the point source which is continuously feeded from outside will result in the condition *∂*_*r*_*u*(0, *t*) = *q*(*t*). As a result, the solution as a function of the radial co-ordinate should have a form *u*(*r, t*) ∝ exp(− *r/σ*(*t*)) in the close vicinity of the distribution’s top.

The completely quantitative test of the applicability of the proposed approach to own new quantitative simulated and experimental results will be given in the next section.

### Experimental data

The experimental data on isolated leakage events after BBB opening were obtained as follows. For intravital experiments, male mice (from 20 to 25 g) were housed under standard laboratory conditions, with access to food and water, *ad libitum*. All procedures were performed in accordance with the “Guide for the Care and Use of Laboratory Animals” [27]. The experimental protocol was approved by the Committee for the Care and Use of Laboratory Animals at Saratov State University (Protocol 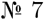, 07.02.2018).

The skull optical clearing method FDISCO was applied to establish an optical clearing skull window [28, 29]. The opening of the blood-brain barrier was performed by sound in accordance with the protocol described in [30]. The animal was placed in a soundproof container together with a sound source (100dB, 37 0 Hz). To detect the Evans blue dye inside the animal’s circulatory system, a fluorescence microscope was used. The images were recorded by a CMOS camera (uc-480, Thorlabs, USA), with a resolution of 1240 × 1080 pixels, at a frame rate of 24 fps. The further equipment details can be found in [29]. The exemplary recorded image is shown in Fig. 1, (A).

**Fig 1.**
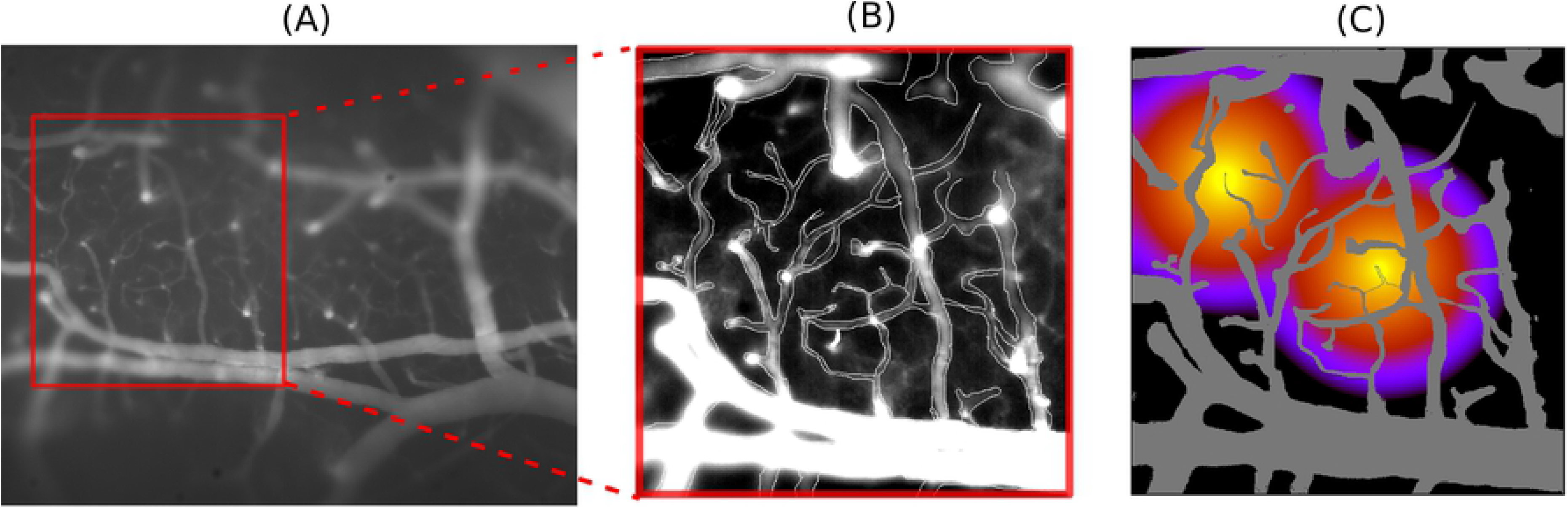
Simulation of diffusion in the realistic template. (A): The flourescent image of the brain vasculature obtained as described in “Experimental data” section; (B): Binarization of selected region; (C): The resulted computation template together with computed results of two-source diffusion. The vessels are painted in uniform grey. Two sources are both non-point, having realistic size in pixels (10-50). Colorbar is from 0.0 (black) to 1.0 (yellow).

Besides the above-described data, we illustrate our method using a few images taken from Ref. [31]. These data published under CC BY 4.0 license and showing 66 kDa fluorescent tracer spread in mouse’s brain parenchyma.

### Simulated data

The experimental data can not be used for the method validation since many parameters are difficult or impossible to evaluate. For this purpose, we used the data obtained by numerical simulation of a diffusion process on a 2D lattice with various kinds of inhomogeneities and taking into account the different size and shape of the source of diffusing substance.

The computer simulation makes it easy to control the process: to include a constant flow of matter into the source, simulating a leaking BBB, or to stop it to simulate a purely diffusive spreading. We assumed that the processes under study have a negligible contribution from advection (the transfer of substance matter with the fluid flow), so it was not included in the mathematical model.

Since we model the image on a discrete pixel lattice, the single-layer version of the mathematical model consists of *N × M* coupled ODEs of the form:

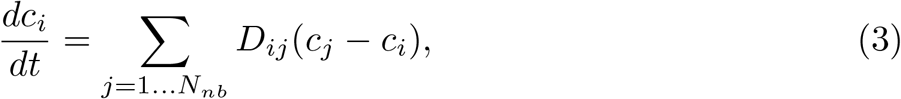

where *N*_*nb*_ neighboring elements contributes to the flows of *c* into and out of *i*-th grid element. Depending on the type of *i*-th and *j*-th elements, the coupling factor *D*_*ij*_ takes the values *D*_1_ if the neighboring pixels both belongs to the intercellular space, and 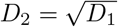 if one or two of them belong to the vessel in order to take into account the slowdown in the spread of the fluorescent dye when flowing around the vessels above and below the observation plane. The process of obtaining a template for calculations is illustrated in Fig.1, where the panel (A) shows the initial fluorescence image (without BBB leaks), the panel (B) illustrates the process of binarization of the vessel structure, and the panel (C) shows the results of the simulation of the process of spreading the substance from two non-point sources. During the testing of the method, we used various combinations of the vessel patterns and sources, from the simplest to the most realistic, as specified below.

## Results

In this section we describe the results of five subsequent trials of increasing realism.

Trial 1 plays the role of a simple test, for which the results are known in advance, so we can demonstrate that method works correctly. Here we analyze simulated diffusion starting from 1 or 2 points of initial nonzero concentration, with no leakage from sources after the initial time. Trial 2 also uses simulated data with small (2×2) single source, but with another leakage scenario. The obtained results argue for the need for additional distribution analysis to select the most appropriate kernel function for blurring. Trial 3 regards to the non-point (like vessel segment) source, with constant leakage and with “shadowing” by nearby located vessels. This is the most realistic simulation of experimental data, which are analyzed in trial 4, where the processing of experimentally recorded local event of dye leakage from opened BBB is described. This trial deals with the unknown in advance leakage dynamics. At last, trial 5 shows the results of the processing of experimental data with multiple sources and unknown in advance leakage scenario.

### Trial 1: Simulated data, point sources

To test the suggested approach, let us consider how does it work for the case of point sources. Fig 2 (A) shows the snapshots of color-coded distributions for the one- and two-point sources, top and bottom row, respectively. In the both cases the source points are not supplied from anywhere, so the amount of spreading substance is preserved. One can see the bell-shaped (overlapped for two-point source) distributions with decreasing maxima. For the single-point source *v*(*x, y*) = *δ*(*x, y*), and the sequence of trial functions (2) reduces to the Gaussian functions

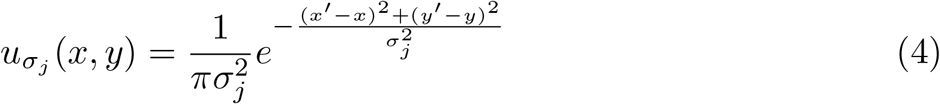

calculated within the interval *σ*_*j*_ = [1, 30] with unit step. The analysed sequence of solutions of Eq (1) has the same functional form

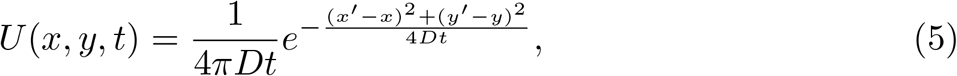

where *D* is the diffusion coefficient, which have to be determined (during this trial we set *D* = 2 a.u.).

**Fig 2.**
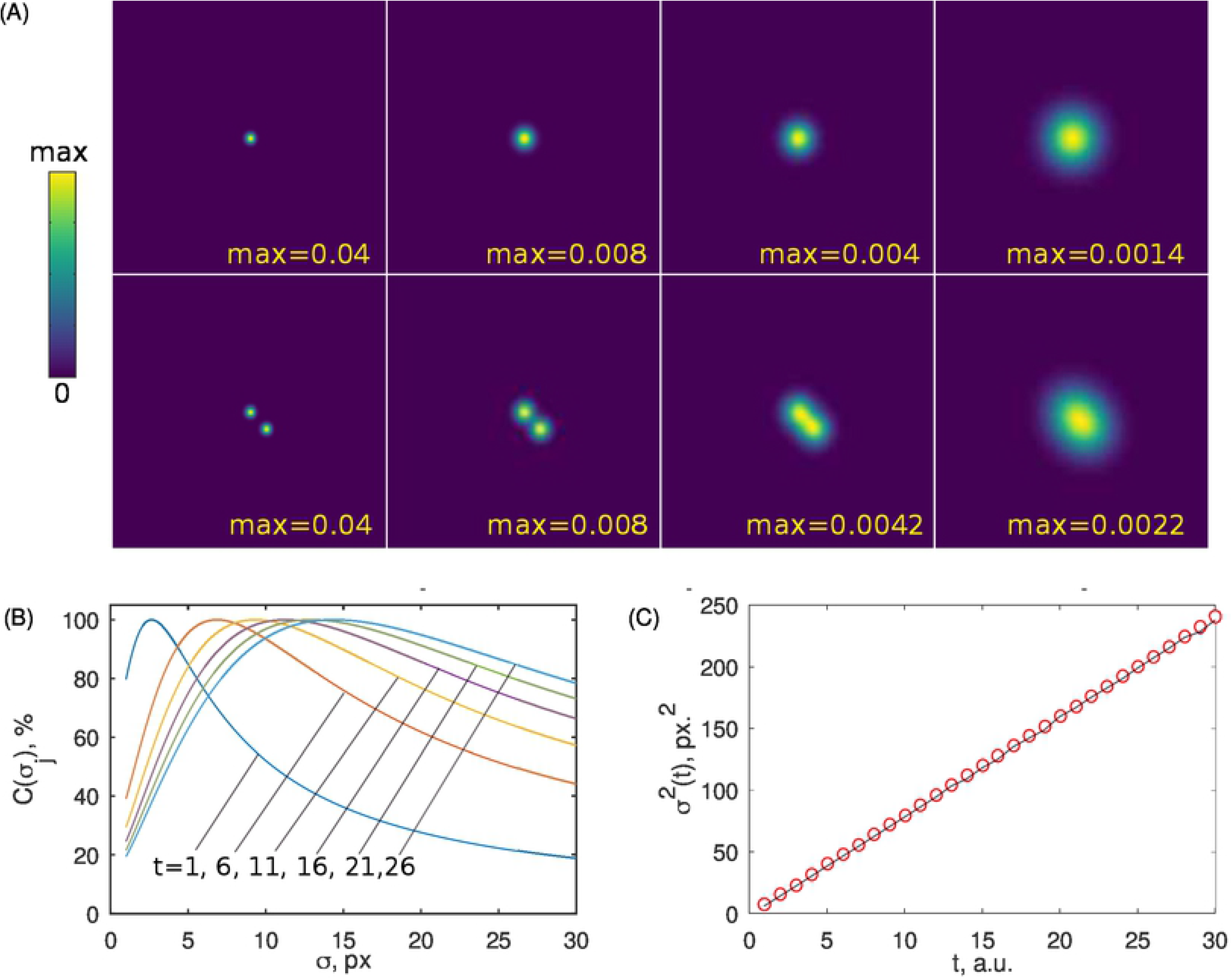
A test of the correlation-based algorithm with the point sources. (A): An illustration of the computed diffusive spreading of a substance from one and two point sources.The upper and the lower rows shows the marker’s density distributions for the time moments *t* = 1/8, 4, 10, 30 a.u.. B: Correlation coefficient between the analyzed images and its counterparts obtained by Gaussian blurring of the initial image versus dispersion *σ*. Curves are drawn for sequentially growing time moments *t* = 1, 6, 11, 16, 21, 26, from the left (blue) to the right (light blue). No difference were found between the curves for one and two sources, so we show only one set of it. C: The dispersion squared for the normal diffusion (solid line) and for its values determined from the maximum correlation (circles) as shown in the panel (B).

Since this initial condition describes a discrete point source we took *σ*_1_ = 1 in Eq (4) that is equivalent to Eq (5) with *t* = 1/8. Fig 2(B) shows the correlation curves between the functions (4) and (5) computed for the sequential time moments started from *t* = 1 a.u. separated by step *δt* = 5 a.u.. As one can expect, the values of their maxima indicate the complete (100%) correlation due to the coincidence of two functional forms. At the same time, the location of the maxima as a function of the dispersion *σ* moves from left to right with the growing time since for pure diffusive process *σ*^2^(*t*) = 4*Dt*. As a result, the dispersion squared grows linearly with time and perfectly coincides with the one, which follows from the analyzed Gaussian spread, see Fig 2(C). The slope of the linear regression line plotted along with the crosses gives the searched diffusion coefficient accurately.

In the case of the two-point source, see the most left picture in the second row of Fig 2 (A), initially, there are two peaks, each concentrated in one pixel, separated by 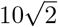px (there are distances in 10 px along each axis). During the time course, two localized spots become wider and even merging into one but not circular shape, see the second row in Fig 2 (A). The correlation-based procedure with initial conditions *v*(*x, y*) = *δ*(*x, y*) + *δ*(*x* − 10, *y* − 10) results in the set of functions 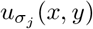, which are linear combinations of the Gaussians. Thus the respective correlation curves and the function *σ*^2^(*t*) are essentially the same as presented for the single-point source(not shown). This confirms, that the diffusion coefficient can be accurately determined and in case of multiple sources as well.

### Trial 2: Simulated data, non-stationary source

This trial intended to argue for the need of the “spreading kernel test” to select the most appropriate distribution function. Since in realistic conditions we do not know the parameters of the leakage, it is not possible to find the exact analytical solution. Instead, we can simply check this functional form by numerical simulations on a lattice.

Here we describe the processing of simulated data using small (2×2 pixels) source, but with time-dependent properties resembling the dye leakage through the opened BBB: source is externally supplied in the limited time interval *t* = 30.0 … 300.0. Thus, during this time interval the total amount of spreading substance increases, while the concentration in the source is maintained constant. Fig 3(A) shows the cross section of the solution. The plot is given in semi-logarithmic axis. During *t* = 30.0 300.0 the distribution profiles are triangle-shaped. Further, the tail part spread faster but the overall shape is still comprised of practically two straight lines. For *t* > 300 the shape of distribution transforms to the parabolic form, which in semi-logarithmic co-ordinates matches the Gaussian distribution.

**Fig 3.**
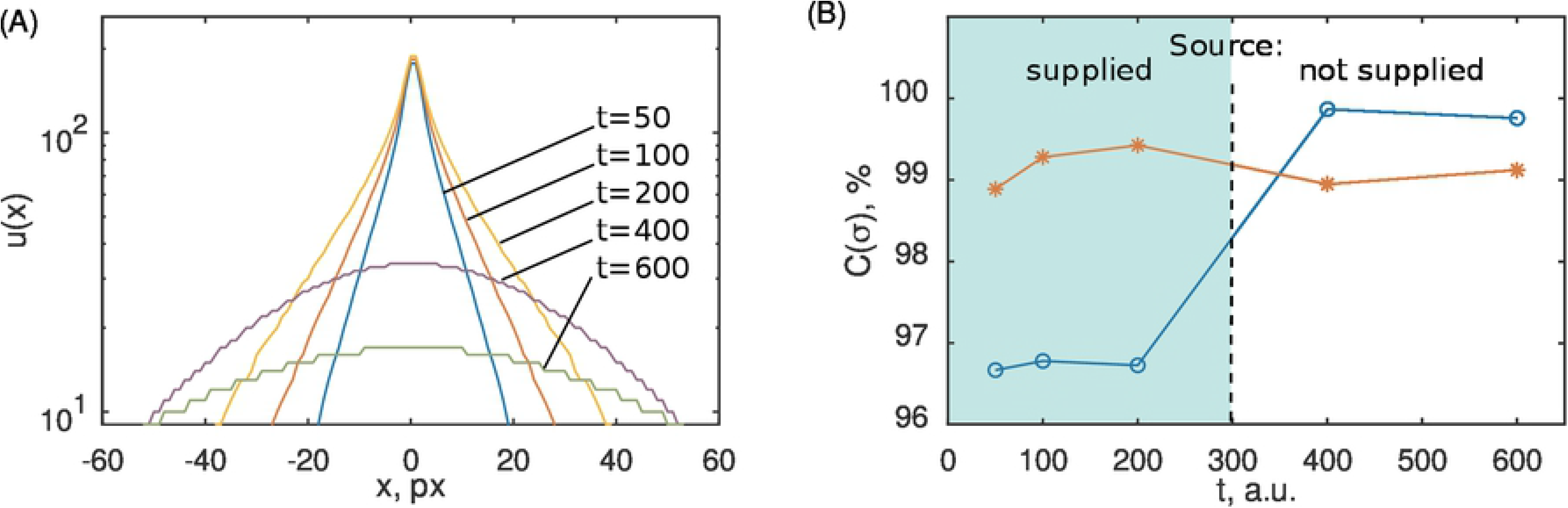
Transitions from exponentially to Gaussian-distributed profiles due to switching off the supply of the source. (A): The central cross-sections of the axially symmetric distributions emerged due to the external supply of the 2 2 pixel source switched off at the time moment *t* = 300 a.u.. Curves are shown for the time moments *t* = 50, 100, 200, 400, 600 a.u.; u(x) is measured in pixel values for 8-bit greyscale color encoding; (B): The correlation coefficients obtained from the convolution-based smoothing with the exponential (red, asterisks) and the Gaussian (blue, open circles) kernels

The above described results demand the improvement of the correlation-based method for estimating the spread parameters, while does not affect its key idea. Namely, if externally supplied source is expected, the set of trial Gaussian kernels (4) should be updated with the set of exponential kernels

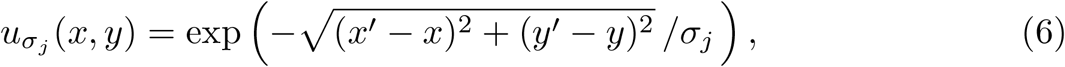

used in the same way as Gaussian ones.

Further, the additional step is introduced being the comparison of the obtained best correlations for these two kernels, as shown in Fig 3(B). One can easily observe the time intervals of the prevalence of each kernel, so we can assess the source dynamics. If the specific purpose is to determine the diffusion coefficient of the parenchyma under conditions of the free diffusion, one should exclude the time interval with the active leakage from the analysis.

### Trial 3: Processing simulated images: non-point and non-stationary source

Now we apply the proposed method to the simulated system taking the region, comprise both the vascular system and the paravascular space. The goal is to determine the accuracy of the diffusion coefficient determination quantitatively. In this section, we use the computational template based on a recorded image as shown in Fig 4, part I (A) and (B). At the initial stage, the visible asymmetry of distribution is due to the source geometry, while at the later stage becomes more symmetric but still imperfect due to the more slow spreading over “vessels”.

**Fig 4.**
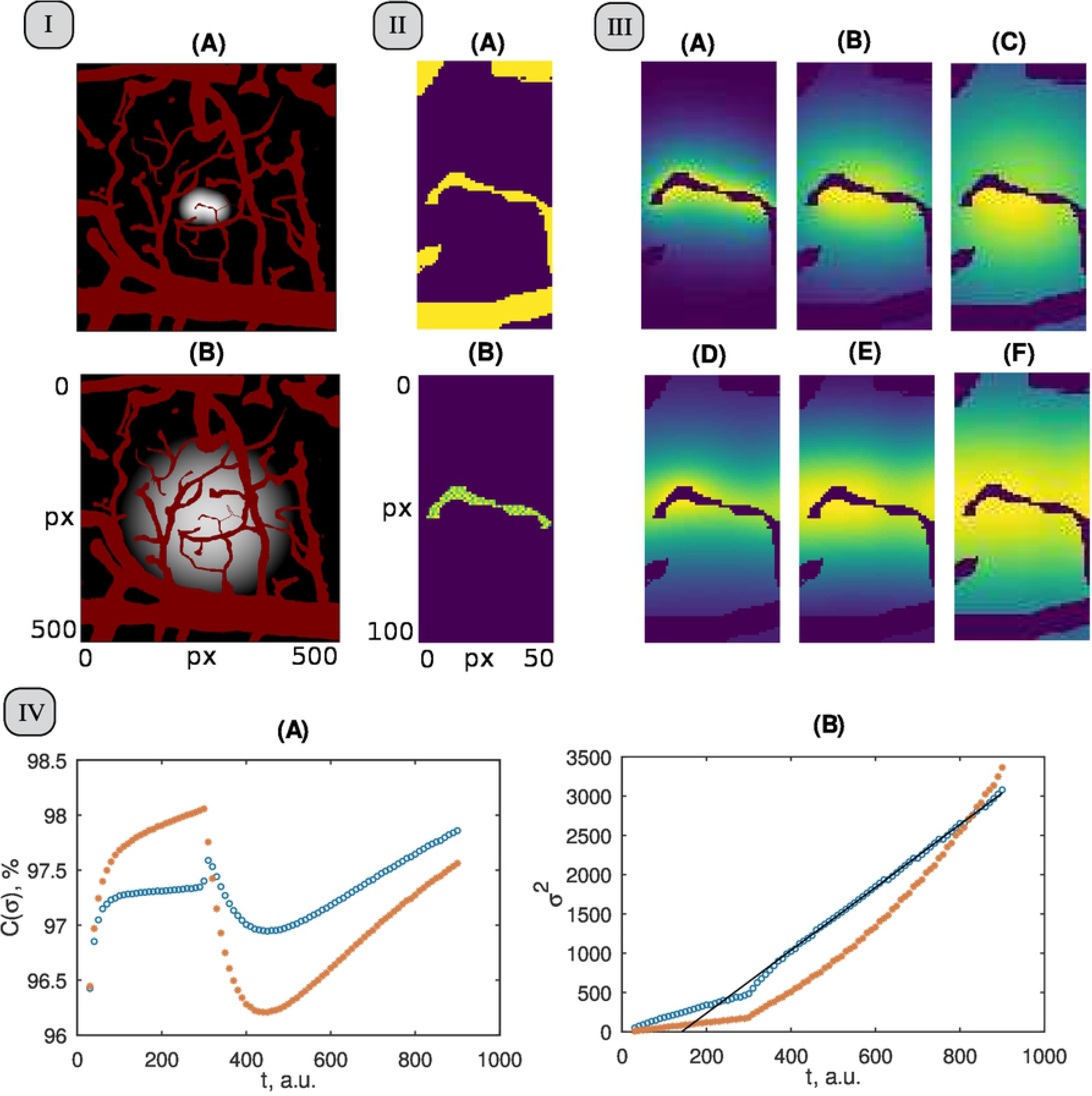
A test of the correlation-based algorithm with the simulated BBB opening event. Part I: Simulated diffusive spreading of a substance using experimental based template at *t* = 50 (A) and *t* = 1000 (B). The logarithmic scale for grayscale bar was used. Part II: The region of interest surrounding the source (A) and the automatically detected source area (B). Part III: Panels (A), (B), and (C) show the simulated distribution of the dye at time moments *t* = 150, 350, and 650, respectively. Panels (D), (E), (F) show the approximation for the same time moments using kernel, showing maximal correlation with the processed images. Specifically, the panel (D) uses the exponential kernel, while (E), (F) use the Gaussian smoothing. Part IV: Time evolution of the maximal correlation coefficients between the simulated and blurred images (A); The growth of the dispersion squared (B). Circles (blue) and asterisks (red) denote results obtained for the Gaussian and exponential smoothing kernels, respectively.

The region of interest was selected in the central part of the simulated area as shown in Fig 4, part II (A) and used as an initial condition for Eq. (2). The source (the “leaking part of the capillary”) was detected automatically using the subtraction of few very first frames, part II (B).

Then, the images blurred using both Gaussian and exponential kernels were counterparted to the simulated frames, as shown in Fig 4, part III. Here, panels (A), (B), (C) show the simulated distribution at *t* = 150, 350, 650, while panels (D), (E), (F) show the best-matched blurred images. Note that the selected part of the full simulated region contains now some area occupied by vessels. To avoid their influence on the correlation coefficients, the signal intensity in corresponding pixels in both pairwise image sequences were excluded from the comparison by filling them with zeros accordingly to the mask suggested by the template used.

Fig 4, part IV demonstrates the curves of maximal correlation and the time dependence for the smoothing kernel’s dispersion squared for them.

Note, that the non-stationarity of the leakage process induced a multi-stage procedure, which complements the pure diffusion smoothing algorithm described above for the case of an initial point source. Specifically, we compare in more details the results provided by the smoothing using both types of kernels, the Gaussian and the exponential ones.

Although maximal correlation values in Fig. 4 part IV, (A) denoted by circles and by asterisks are sufficiently high for the whole time region from *t* = 30 a.u. to *t* = 900 a.u., one can see the drastic difference in their behaviour on the time sub-intervals before and after *t* = 300 a.u. that also seen in the dynamics of the dispersions shown in Fig. 4 part IV(B).

For *t* < 300 a.u. (the former case) the smoothing with the exponential kernel results in a higher correlation than with the Gaussian kernel. For *t* > 300 a.u. the correlation curves are interchanged.

Respectively, the circles in Fig. 4 part IV, (B) which goes to 300 a.u. demonstrate some kind of slowly growing function *σ*^2^(*t*). Note that it is concave up that also indicates that the underlying process is questionable as the normal diffusion.

This regime instantly stops at the time moment *t* = 300 a.u. that corresponds to the moment of the leakage’s stop, when the correlation coefficients are shown in Fig. 4 part IV, (A) demonstrate a discontinuity with the jump to a higher value. Further, up to approximately *t* = 400 a.u., there is still non-linear dependence of the dispersion squared, which however gradually tends to the straight line tending further up to the end of recordings. Its linearity is highlighted by the black solid line whose coefficients are obtained with the least-square fitting method for *t* ∈ [400, 900] a.u. sampled with the time step *t* = 10 a.u.. Remarkably, the obtained slope coefficient is equal to 4.0 (with the absolute deviation 0.005) that exactly matches the free diffusion coefficient used in simulations.

The use of exponential kernels (6) instead of the Gaussians (4) in Eq (2), see asterisks Fig. 4 part III, (B), demonstrates a kind of parabolic function there, i.e. the absence of a constant parameter of exponential smoothing with respect to the second moment of the intensity distribution in contrast to the first stage of the process.

Thus, we can conclude that the combination of the plots of the correlation coefficients and the *σ*^2^(*t*) curves allow for distinguishing between the presence and the absence of a supply of the source. As well, in the latter case, it is possible to determine accurately the diffusion coefficient within the localized time interval extracted via the discussed criteria.

### Trial 4: Processing experimental images of isolated BBB leakage event

The experimental grayscale images of the isolated BBB opening event were evaluated as follows. The image taken at 42 s after the beginning of registration was chosen as the initial one since it shows a certain small leakage from a capillary in one of its part. The image was cropped around this capillary, see Fig 5, part I, panel (A).

**Fig 5.**
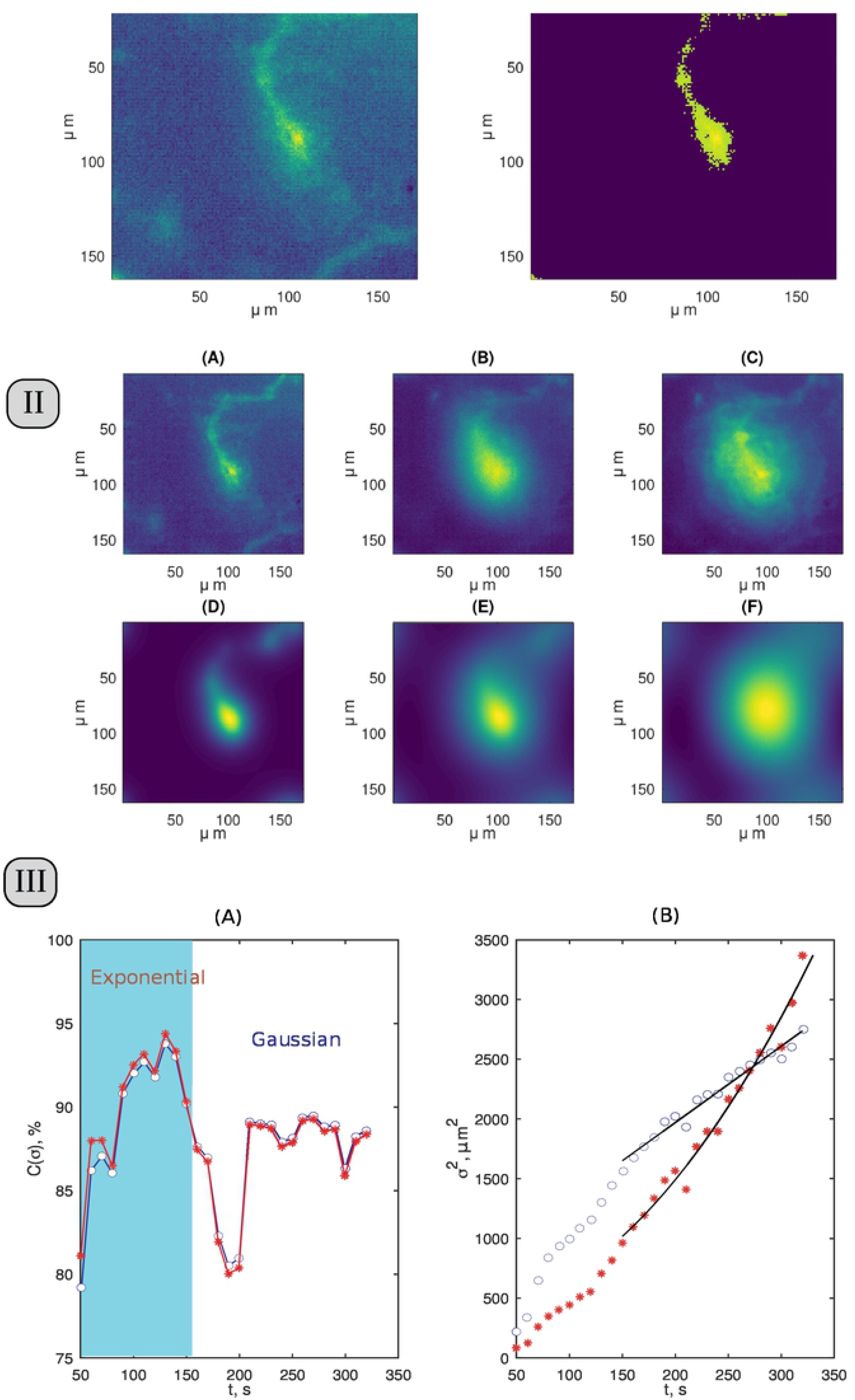
Processing data on isolated BBB leakage event. Part I: (A): selected ROI at *t* = 42. (B): Binarized image of ROI as the template for smoothing. Part II: The comparison of experimental fluorescent marker spreads and the calculated diffusion spreads for the same time moments. The time moments are *t* = 50 s (A) and (D), *t* = 120 s (B) and (E), and *t* = 200 s (C) and (F). Part III: The plot of 2D correlation coefficients and dispersion squared between matrices. Panel (A): the correlation coefficient as a function of the marker spread time, where circles and asterisks mark the usage of the Gaussian and the exponential kernel, respectively; (B): the model dispersions squared calculated using the Gaussian kernel dispersion for the time course of the spread process (circles); the solid line shows the linear fit for the pure diffusive spread regime.

The source configuration was estimated from this image by means of unit matrix, whose elements are equal to 1, when the corresponding element value of the image matrix is larger than 30% of the maximal one, and 0 otherwise. This matrix was used as a mask (i.e. multiplied element-by-element) for the initial image , and converted to floating-point numbers of double precision. This delivered the estimated image of the source *v*(*x, y*) as shown in Fig 5, part I, panel(B).

This initial distribution was substituted into Eq. (2) and the blurred images were obtained for the set of trial *σ*_*j*_.

Fig 5, part II illustrates the comparison of the experimentally registered images around the vessel with leakage and the model spread distributions, where Figs 5, part II, panel(D) and (E) are obtained using the exponential kernel for the smoothing of the initial source shown in Fig 5, part I, panel(B), and Figs 5, part II, panel (F) is the blurred image.

Similar to previous trials, the two-dimensional correlation coefficient *C*(*σ*_*m*_) was calculated with the set of images registered with 10 s interval; the values of providing the best similarities are shown in Fig 5, part III, panel(A) with circles and asterisks for Gaussian and exponential kernels, respectively.

One can see that, except the very first point, the correlation coefficient varies from 80 % to 95 % that can be considered as a pretty high value taking into account certain noise, inhomogeneities in the experimental recordings, and simplicity of the model. For the *t* < 150 s the exponential kernel provides better correlations in comparison with the Gaussian one. However, this advantage in the correlation values is not high.

As well, Fig 5, part III, panel(B) exhibits the courses of the function *σ*^2^(*t*) for both kernels.

Comparing with previous trial, we can draw a certain analogy. Namely, the initial phase of the process shows its own dynamics, in which what happens to the source, and not the distribution of matter, plays the main role. It is difficult to expect that the actual process being analyzed upon opening the BBB will follow a simple scenario of trial 3. It is more likely that the outflow of dye from the capillary follows an irregular pattern. This may explain the absence of the drastic prevalence of the exponential kernel-based correlations even during the first stage of the process.

Later, the process goes to the stage where the main contribution is made by the spreading of substance through diffusion. For this second phase, the curves for *σ*^2^(*t*) are very similar for trial 3 and trial 4. The curve for Gaussian smoothing looks quite similar to the course of the same function in Fig 4, part III, panel(B) for (*t* > 300 a.u.). i.e. non-linear growth followed by the diffusive spread without the constantly acting source. Note that for both trial 3 and trial 4, in this phase, the graph *σ*^2^(*t*) for the exponential kernel shows increasing steepness.

Now, we can quantify the process revealed as the normal diffusive spread that characterizes the properties of the brain’s parenchyma. In Fig 5, part III, panel(B), the respective region corresponds to *t* ≥ 140 s, i.e. within the interval, where the Gaussian-smoothed model gives the better correlation than the exponential kernel-based one. Straight black line in Fig 5, part III, panel (B) shows the fitting result using the expression 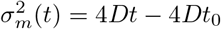 in order to evaluate the *D* value. The correlation coefficient between the shown line and the data is equal to 98.5 %. The diffusion coefficient determined from the slope of this straight line is equal to *D* = 1.59 (*μ*m)^2^/s; its interval varies from *D*_−_ = 1.45 (*μ*m)^2^/s to *D*_+_ = 1.74 (*μ*m)^2^/s with 95 % confidence bounds. The quantitative relevance of obtained *D* assessement we further discuss in conclusion section.

### Trial 5: Processing experimental images with multiple emerging sources

Finally, we present the challenging trial which runs beyond the capability of the proposed method in its present form. Specifically, we analyze the experimental data presented in [31] (published under CC BY 4.0 license). The sequence of the four images presented in Fig 6 (A)-(D) illustrates the massive leakage of 66 kDa fluorescent tracer from multiple sources distributed over the whole cortex surface. The visual inspection of the images indicates that the tree-like geometry of the sources evolves in time alongside with tracer spreading. Physiologically, it is likely due to the gradual penetration of the tracer into the perivascular spaces of the more and more small vessels, compare the regions indicated by the arrow in the images (A) and (B).

**Fig 6.**
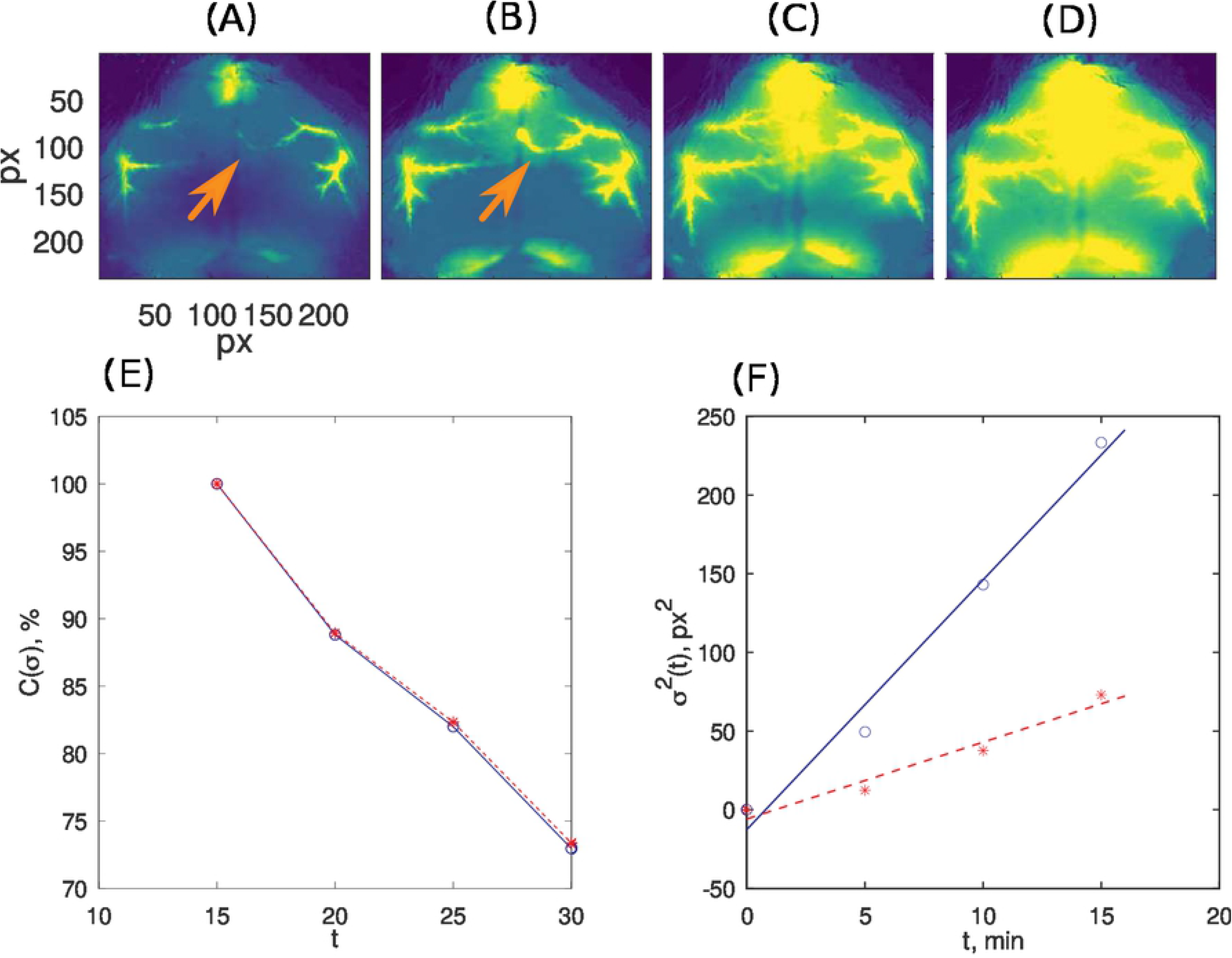
The spreading of the fluorescent tracer from multiple and evolving sources. The upper row of images represents processed experimental images from [31] (published under CC BY 4.0 license). Panel (E) shows the maximal achieved correlation using both the Gaussian and the exponential blurring kernels The panel (F) demonstrates the time evolution of the revealed dispersions squared and its linear fit.

Clearly, such a time-evolving source does not fit the assumptions that underlie our method. However, we still apply it in order to make a kind of “crash test”, to learn how specifically the method will fail. We use the image (A) as a template to determine the source area *v*(*x, y*) for Eq (2).

The panel (E) shows the maximal correlation between experimental and diffusion trial images for Gaussian and exponential blurring kernels for the three moments of time, 5, 10, and 15 min, respectively. They rapidly fall from image to image for both smoothing kernels, showing the increasing inconsistency between the actual process and approximations. Note, the plots for dispersion squared shown in panel (F) still lie close to the straight line, but the relevance of its slope is very questionable in this case.

## Conclusions

Summarizing, in our work, we proposed a simple method to quantitatively characterize the process of spreading of a fluorescent dye over a series of images recorded in an experiment. The proposed method is based on the very basic features of diffusion but does not rely on the assumption of a point source, although it is applicable in this simplest case. The method allows one to assess the evolution of the process. Specifically, we demonstrated how to distinguish, between the constantly feeded and unfeeded sources.

We also believe that our method is less sensitive to the heterogeneity of the medium since it is based on an analysis of the entire luminosity landscape. Thus, the specific choice of a point for analysis is not a challenge, since there is no such choice itself. For the above reasons, it becomes possible to compare the results obtained on different samples.

Our analysis of both surrogate and experimental data on the isolated BBB opening event shows that the correct estimation of the initial moment of time when the dye starts to leak, is unlikely can be made using late-time distributions. Instead, one should find the transition between (at least) two process phases in order to search for the initial phase of the process and calculate the moment of the beginning of the “leak” from it. The proposed method allows one to see and evaluate such changes in the dependence of dispersion on time for short times and, most importantly, in what kind of blurring kernel core (Gaussian or exponential) describes the process better.

We showed that for surrogate (computed) data the actual diffusion coefficient is recovered pretty well. However, the exemplary quantitative estimation of the brain parenchymal diffusion coefficient for the Evans Blue dye delivered *D* ≈ 1.6 (*μ*m)^2^/s. Since the Evans Blue dye is highly aggregated with albumin, from previously reported data [32] one can expect 6 times higher *D*, so there are the issues to discuss. One of the possible reasons is the 3D effects: some part of the dye spreads up and down the focal plane and thus becomes highly blurred in the image. Another issue is that our method estimates the late stage of diffusion at “far-field”, where the diffusion coefficient can fall for the highly inhomogeneous medium [33]. We, however, postpone the detailed analysis of quantitative aspects of measurements *in vivo* for future work, since many additional factors should be controlled during experiments to make valuable conclusions.

Undoubtedly, the application of our method involves the processing of a significantly larger amount of data compared with point measurements. However, these data are generally available using modern imaging techniques. We showed that the presence of images of blood vessels in the analyzed region does not fundamentally affect and does not limit the application of the method, although this needs additional evaluation.

## Acknowledgments

We thank Alexander Khorovodov and Maria Klimova for preparation of animals for the experiments.

